# Quantification of rose rosette emaravirus (RRV) titers in eriophyoid mites: insights into viral dynamics and vector competency

**DOI:** 10.1101/2024.05.23.595398

**Authors:** Tobiasz Druciarek, Alejandro Rojas, Ioannis Tzanetakis

**Affiliations:** Department of Entomology and Plant Pathology, University of Arkansas, Fayetteville, AR, USA; Department of Plant Protection, Warsaw University of Life Sciences, Warsaw, Poland; Department of Plant, Soil and Microbial Sciences, Michigan State University, East Lansing, MI, USA

**Keywords:** *Emaravirus rosae*, *Phyllocoptes fructiphilus*, *Phyllocoptes adalius*, interactome, virus titration

## Abstract

Understanding the interaction between rose rosette emaravirus (RRV) and its vectors is pivotal in addressing the epidemic outbreak of rose rosette disease. This study employed quantitative real-time RT-PCR to assess RRV genome copy numbers in *Phyllocoptes fructiphilus* and *P. adalius*, providing insights into the viral dynamics and vector competency. Our findings suggest active virus replication within *P. fructiphilus*, a confirmed vector species, unlike *P. adalius*, highlighting its non-vector status. Furthermore, the study highlights the variability in virus concentration in mites over time, underlining possible developmental stage-specific response and influence of mite lifestyle on RRV retention and replication. This research is the first step in understanding the virus-mite interactome, which is essential for developing effective management strategies against rose rosette disease.

## INTRODUCTION

Eriophyoid mites (phylum Arthropoda; class Arachnida) are the smallest arthropod virus vectors and cause significant losses in food, tree and ornamental crops worldwide [1,2]. Approximately 5000 species of eriophyoids have been described, but the actual number of these mites is hypothesized to be significantly greater [3,4]. As of 2024, eriophyoid mites are verified or suspected vectors of ∼40 plant viruses [5–8]; however, in the metagenomics era, the rate of identifying vectors is not keeping up with the increasing number of virus discoveries [9,10]. There is an even greater knowledge gap in understanding the dissemination mechanisms of eriophyoid-transmitted viruses [6].

The negative-sense, single-stranded RNA (-ssRNA) genus *Emaravirus* (family *Fimoviridae*; order *Bunyavirales*) is an emerging group of eriophyoid-transmitted viruses comprising more than 30 classified and putative species with worldwide distribution and economic impact [2,11]. *Emaravirus rosae* (member: rose rosette emaravirus, RRV) is considered one of the most economically significant emaraviruses, as infected plants die within two to five years after the onset of symptoms [5], affecting the profitability and sustainability of commercial operations and landscapers in the United States [7,12].

RRV is vectored by *Phyllocoptes fructiphilus* Keifer [13] and the recently identified *Phyllocoptes arcani* Druciarek, Lewandowski & Tzanetakis [14,7]. It remains unclear whether the virions are transiently and reversibly retained or if they circulate and replicate within the mite’s body. This study tested the hypothesis of RRV replication in the mite body by assessing the genome copy numbers in a vector (*P. fructiphilus*) and a non-vector (*P. adalius*). This research provides a deeper understanding of the molecular interactions between RRV and mites and offers new perspectives on the factors influencing the dissemination of RRV.

## METHODS

### Maintenance of mites and plants

The avirulent *P. adalius* and *P. fructiphilus* colonies used previously [7] were maintained on potted KnockOut® roses (*Rosa* × *hybrida* ‘Radrazz’) and tested as described previously [13]. RRV was maintained on infected KnockOut® roses by *P. fructiphilus-*mediated transmission. The RRV isolate obtained from these plants was Sanger-sequenced and matched isolates available in NCBI. Mite colonies and RRV-source plants were maintained in separate environmental growth chambers (14L:10D, 20°C, 70% RH) and monitored for several months before being used in experiments.

### Construction of standard curves

Standard curves were generated for each target to determine the absolute number of RRV genome copies in mites. The emaravirus-specific primer PDA213 [15] was used for reverse transcription (RT), generating cDNA from viruliferous *P. fructiphilus* specimen as described below. An amplicon encompassing the virus target region was generated, whereas, for an internal control/reference gene, an amplicon targeting the 18S rDNA region of the mite was also obtained (supplemental material). DNA concentrations of sequenced amplicons were determined with a Qubit 3.0 fluorometer (Life Technologies), and the copy number of each target was calculated using the formula: *V*_*c*_= (*C*_*a*_ x *N*_*A*_)/(*l*_*a*_ x *m*_*b*_), where *V*_*c*_ is the number of virus copies/μL, *C*_*a*_ is the amplicon concentration in ng, *N*_*A*_ is the Avogadro’s constant (6.02 × 10_23_), *l*_*a*_ is the amplicon length in base pairs, and *m*_*b*_ is the molecular mass of 1 bp in ng/mol (660 × 10_9_). Tenfold dilutions (10_6_-10_2_ copies) were prepared, and RT-qPCR was performed with two technical replicates, as described below. Curves were constructed by plotting the *quantification cycle* (C_q_) values versus the log10 of the target copy number. The amplification efficiency (E) of each assay was calculated using the equation *E* = 10^(− 1/S)^, where *S* is the slope of the corresponding curve.

### Quantification of RRV titer

Quantification of viral and reference gene copies was performed using a modified version of the direct RT-PCR method described previously [6] with standards and cDNAs from mite and plant samples assayed by qPCR (supplemental material). Samples were analyzed in two technical replicates for RRV RNA3 and mite 18S rDNA. No-template controls, RRV-free rose, and non-viruliferous *P. fructiphilus* mites were included in the experiments to assess contamination and specificity, respectively. C_q_ values from RRV-containing samples were compared with standard curves to determine the absolute quantities of the targets, with the values normalized by quantities of the corresponding reference gene.

### RRV titer in mites over time

Immature mites (larvae) from each avirulent colony were transferred to modified Munger cells (60/cell) [16] containing detached, RRV-infected leaflets and kept for 24 hours in cells placed in an environmental growth chamber (14L:10D, 27°C, 63% RH) for virus acquisition. There were eight cells for *P. adalius* and 12 for *P. fructiphilus*. On the second day, two mites from each cell were transferred to tubes containing TE buffer and stored at -80°C for subsequent analysis. The remaining mites were subsequently moved to a new cell with a detached, RRV-free leaflet for 24 hours. This process of collecting two individuals and transferring the remaining mites to a new cell with a detached, RRV-free leaflet continued daily until day 8 (Fig. 1). Consequently, 16 mites per day were collected and analyzed for *P. adalius*, and 24 per day were collected and analyzed for *P. fructiphilus*. Additionally, 16 and 24 mites, respectively, were collected from the mite stock colonies just before their initial transfer to the RRV-infected leaflets for virus acquisition. We collected and analyzed 144 *P. adalius* and 216 *P. fructiphilus* individuals throughout the experiment.

**Fig. 1.**
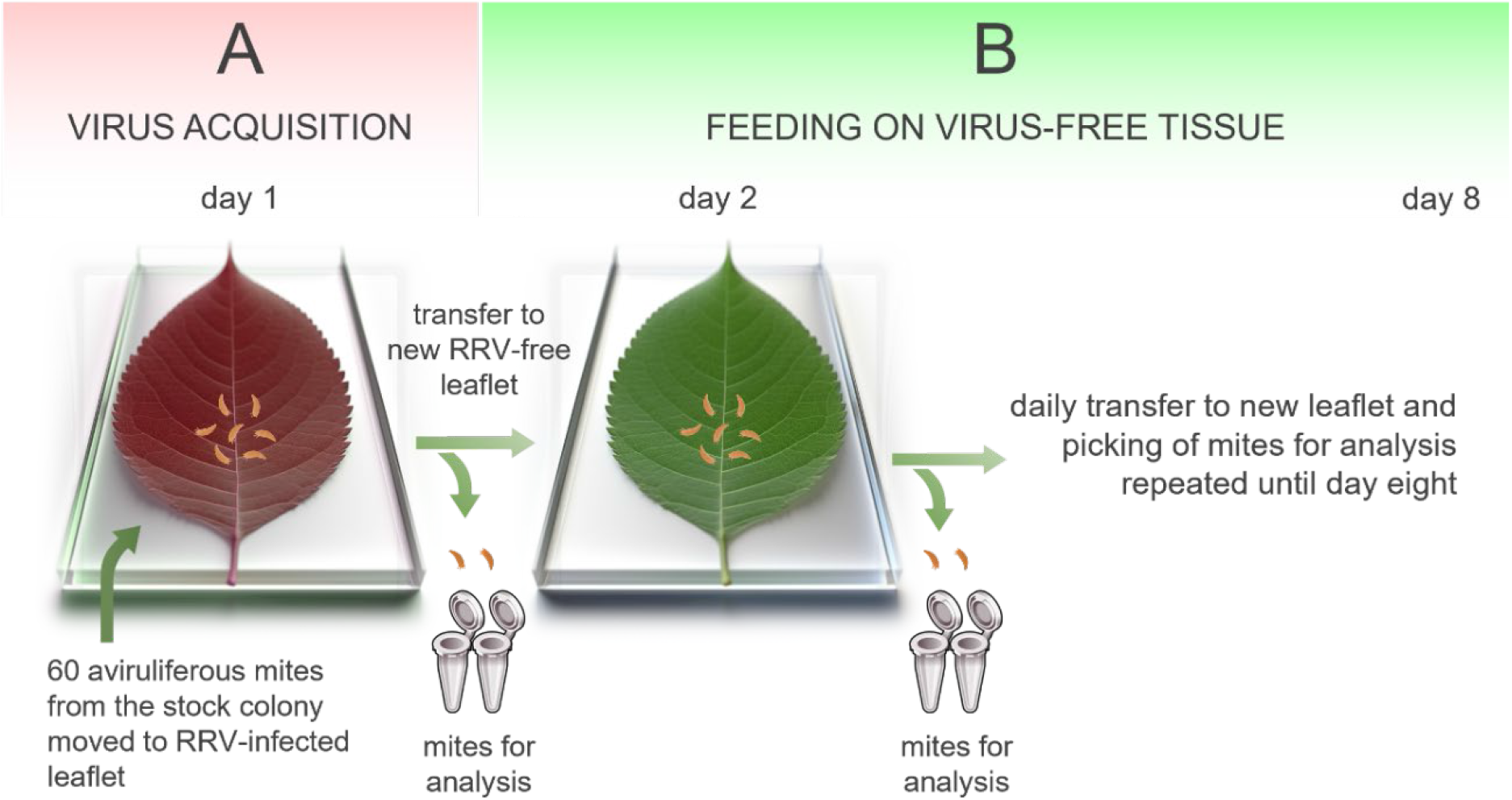
Schematic representation of rose rosette emaravirus (RRV) quantification assay. **A**, Virus acquisition by immature mites moved to RRV-infected material and fed for 24 hours. **B**, Daily transfer of developing mites to new, RRV-free tissue with two mites taken daily for analysis. The artwork was partially produced using the Midjourney bot via a Discord server at https://discord.com/invite/midjourney

### Statistical analyses

The resulting qPCR runs were extracted using batch processing mode in CFX Maestro v2.3 (Bio-Rad, Hercules, CA) and imported into R version 4.2.1 (R Core Team, Vienna, Austria). Since there are multiple independent qPCR runs, tenfold standards (10_6_-10_2_ copies) were included on every plate for RRV and mite rDNA. The data was analyzed to determine whether there were differences between plates before combining the data for further analysis. A linear model was employed, using C_t_ values as the response variable and log-transformed copies as a factor while treating the plate as a random factor. This approach was used to assess variability across plates before merging the results for comprehensive analysis.

For the merged data, an infection coefficient (IC) was calculated as follows: IC = RRV/mite rDNA concentration. An additional approach to assess infection efficacy was to use a normalized Infection Coefficient (nIC), defined as dividing the C_t_ value of the vector by the C_t_ value for the cDNA of RRV (nIC = C_t_ mite/C_t_ RRV).

Linear regression analysis was performed to assess the concentration of virus in each mite species. Total DNA was quantified from the host mites via qPCR, and the results were compared with the corresponding virus concentrations estimated via RT-qPCR. A constant was added to all virus samples to adjust for zero values, and DNA concentrations for viruses and mites were log10 transformed. A Pearson correlation was calculated to determine whether there was a significant correlation between the two variables. To investigate the differences in the infection coefficient or virus concentration across the eight feeding events (days), the infection coefficient was analyzed over eight days (events). A repeated measures analysis was performed to identify any differences across these events. Both the mite species and the acquisition events were treated as factors in a two-way ANOVA for repeated measures. Significant effects were further evaluated using post-hoc tests, specifically pairwise comparisons with adjustments using the Bonferroni method for multiple comparisons. All analyses were conducted using R version 4.2.

## RESULTS

### Infection coefficient

The factor corresponding to the independent plate was included as a random factor in the analysis, explaining only 0.013 and 0.015 of the variances in the virus and mite rDNA concentrations, respectively. Additionally, the homogeneity of the regression slopes across both assays was tested and found to be statistically insignificant (RRV p= 0.328, mite rDNA p= 0.808) (S. Fig. 1).

Analysis of the change in virus concentration in response to the rDNA concentration of each of the two mite species revealed that for RRV - *P. adalius* rDNA concentration had a statistically insignificant regression (R^2^=0.045, p= 0.29), suggesting that the virus did not replicate in the mites (Fig. 2). In contrast, a positive correlation was observed between RRV and *P. fructiphilus* rDNA (R^2^=0.36, p= 2.2e^-16^; Fig. 2), indicating that the virus concentration increases as it replicates.

**Fig. 2.**
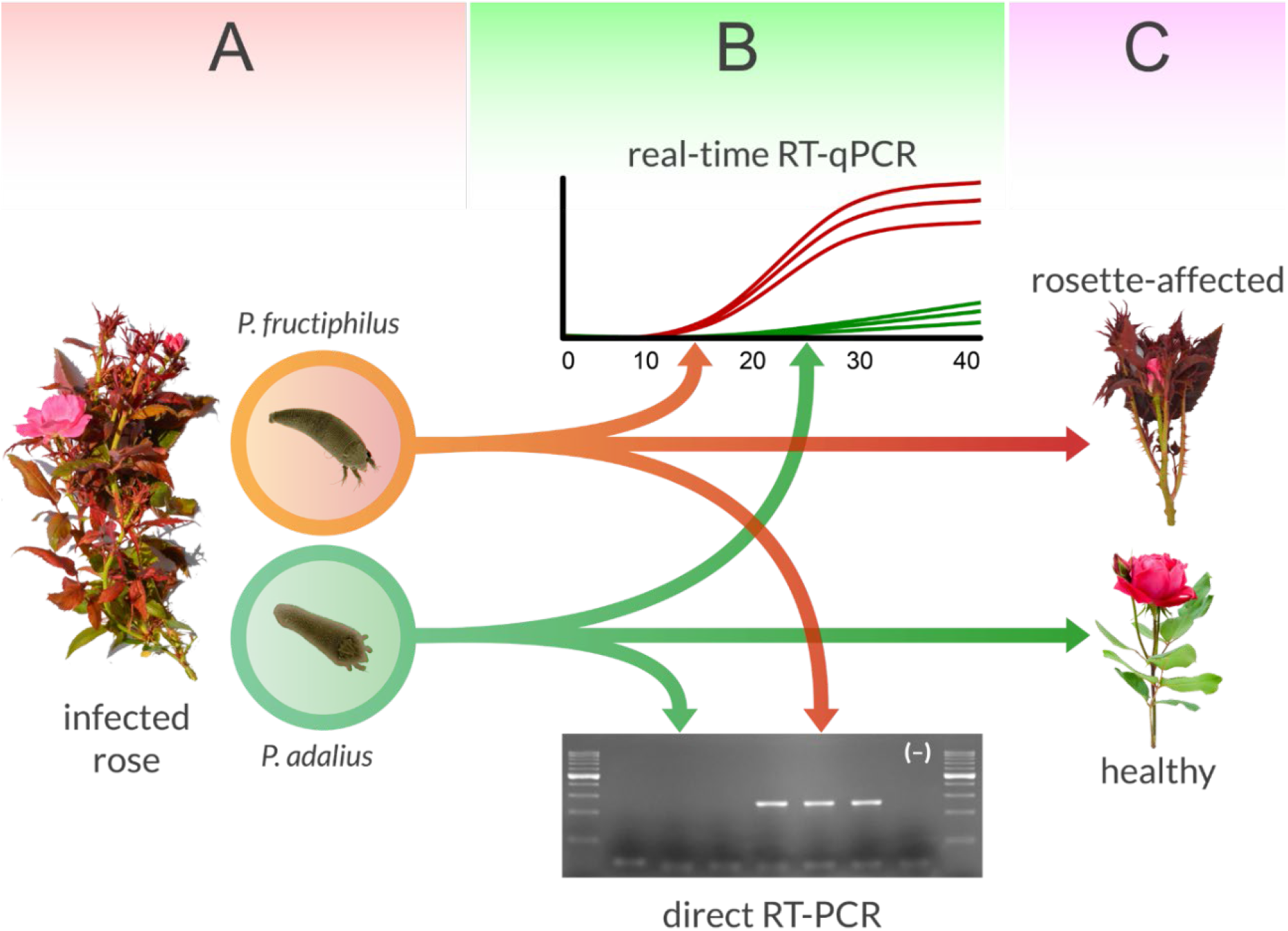
Schematic representation of rose rosette emaravirus (RRV) transmission competency by eriophyoid mites. **A**, RRV might be acquired by both *Phyllocoptes* species feeding on infected rose plants. **B**, However, only *P. fructiphilus* has enough of a virus load to obtain a positive amplicon in semi-quantitative RT–PCR [6], and the RT–qPCR assay suggested replication in this species. **C**, Transfer of viruliferous mites to recipient plants results in successful transmission and development of symptoms only in the case of *P. fructiphilus* [7]

The normalized infection coefficient showed that both species acquired RRV (Fig. 3). The overall infection coefficient varied between 0.3 and 0.6, with *P. adalius* displaying greater variability. Most feeding events yielded similar results; however, on day 5, the infection coefficient for *P. fructiphilus* surpassed that for *P. adalius*. Repeated measures analysis of these fluctuations indicated significant differences in the infection coefficient at days 0, 1, 2, 5, and 8 (Fig. 4). In particular, *P. adalius* had higher coefficients on days 1 and 2, although the difference was less significant than that in the instances where *P. fructiphilus* dominated (days 0, 5, and 8). While the trend was consistent for the initial events, day 5 marked a notable increase (p= 1.35e^-14^) in the virus concentration for *P. fructiphilus*.

**Fig. 3.**
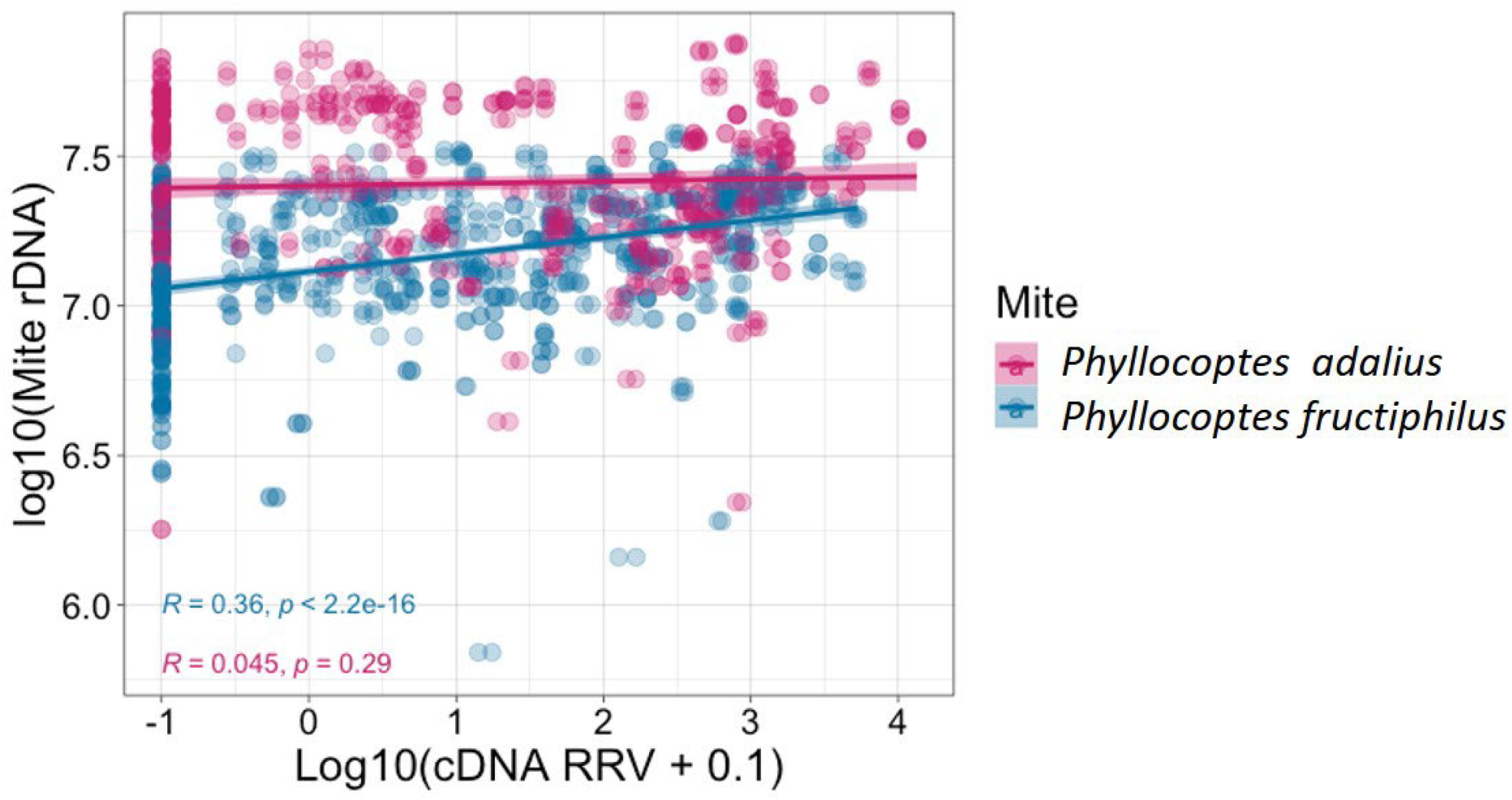
Correlation of virus load (log10 of cDNA ng/µL) and the corresponding host mite DNA (log10 of mite rDNA). The bands represent the 95% confidence intervals of the fit lines. Correlations were evaluated using Pearson correlation, and R-square and p-values for *Phyllocoptes adalius* and *P. fructiphilus* were included

**Fig. 4.**
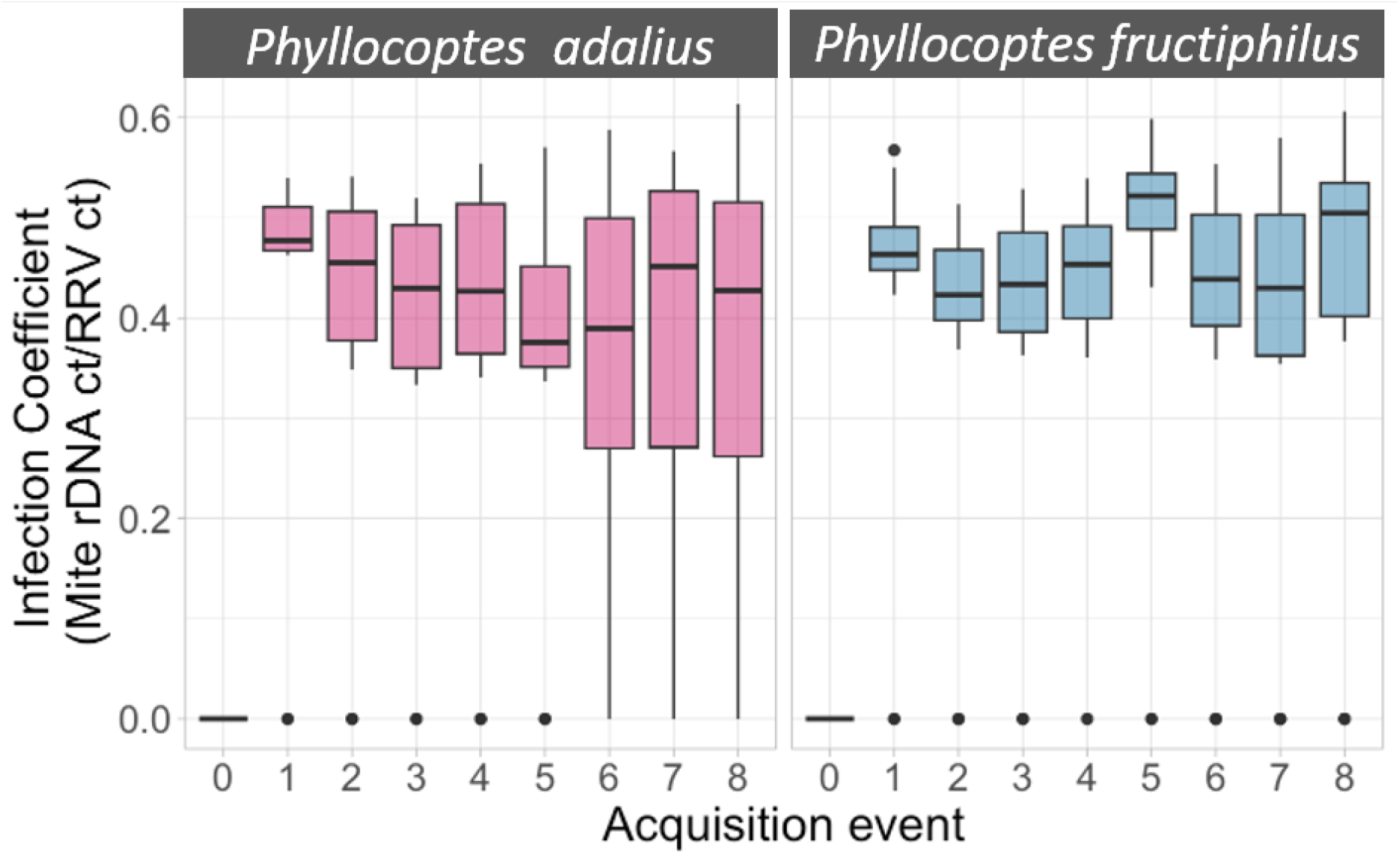
Box plot of the normalized infection coefficient of rose rosette emaravirus to *Phyllocoptes adalius* and *P. fructiphilus* per acquisition event. Dots represent outliers

## DISCUSSION

Our study advances the understanding of virus dynamics by quantitatively monitoring virus concentrations over time in mites transiently exposed to RRV-infected tissue. We cleared the digestive tract and prevented further uptake of infected plant material by transferring mites to virus-free tissues daily and quantifying the viral concentrations. The use of *P. adalius*, a non-vector species, and *P. fructiphilus*, a verified RRV vector, provided a new perspective on vector competency and virus-mite interaction dynamics (Fig. 2) [6,7,13].

The quantitative assay enabled RRV and mite rDNA assessment, revealing acquisition by both species (Figs. 3 and 4). The infection coefficient, derived from RRV/rDNA concentrations and C_t_ value ratios, revealed new aspects of RRV dynamics. Notably, there was a positive correlation between the virus concentration and the vector rDNA concentration in *P. fructiphilus*; as the number of rDNA copies increased (presumably, immature mites develop into adults), as did the virus concentration within the mite, indicating replication of RRV in a verified vector. These results agree with those reported previously [6], in which amplicons were obtained from *P. fructiphilus* but not *P. adalius* individuals.

The variability in the infection coefficient, especially the spike in *P. fructiphilus* on day 5, suggests factors influencing RRV dynamics at different mite developmental stages (Fig. 5). Interestingly, on day 5, RRV transmission was also reported previously [13]. Considering the developmental times for life stages previously reported for both species [16,17], it is highly probable that by day 5, mites had reached an adult stage. We initiated the study with cohorts of immature individuals to ensure virus acquisition and that enough individuals were alive throughout the experiment. However, our methodology also had several limitations, as it prevented us from verifying the specific life stage at sampling, which could have provided detailed insights into the stage-specific virus response.

**Fig. 5.**
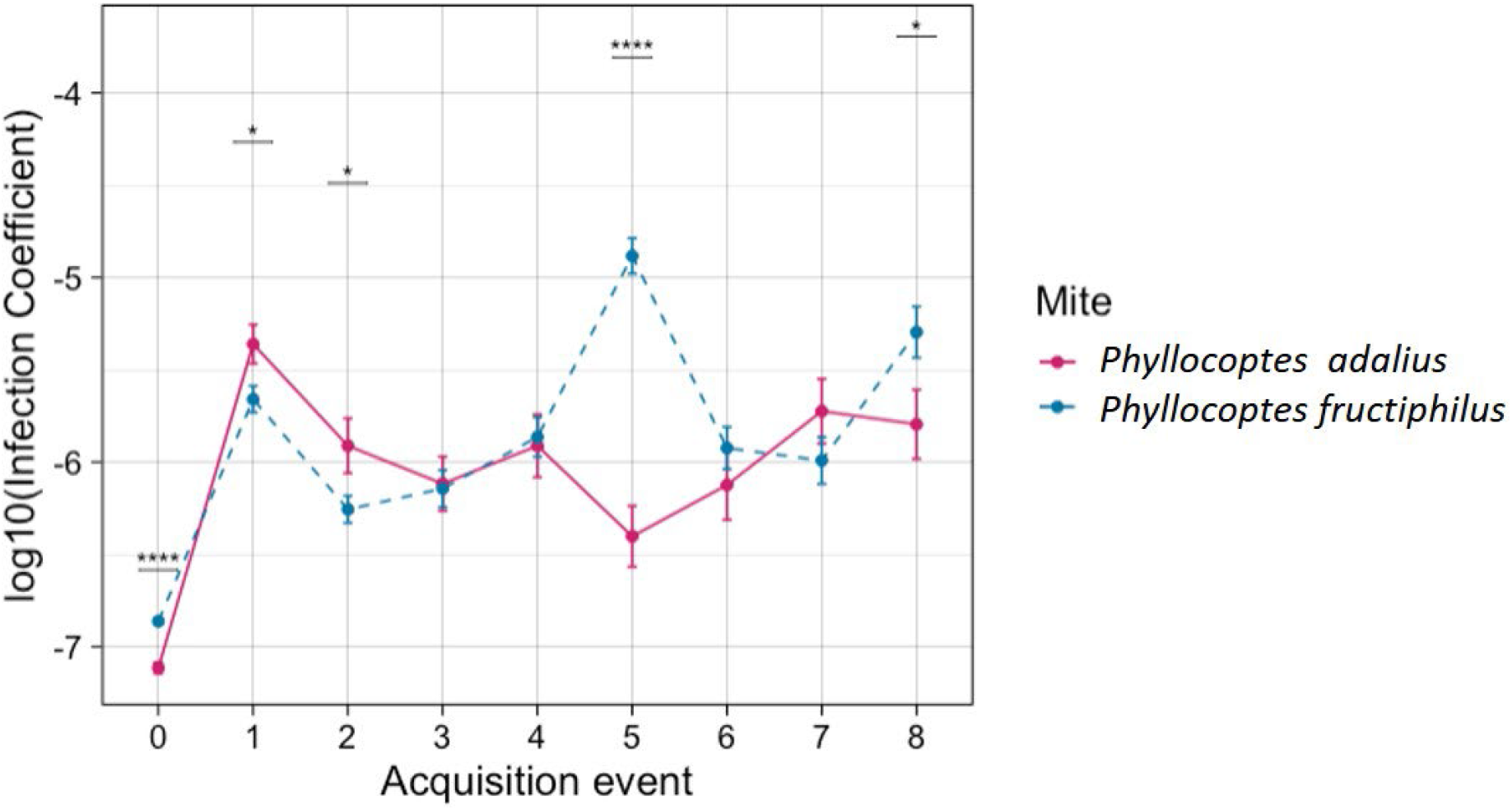
Acquisition event dynamics of rose rosette emaravirus (RRV) log10 infection coefficient derived from the cDNA RRV divided by the mite rDNA. Points represent the means of 16 and 24 individual mites for *Phyllocoptes adalius* and *P. fructiphilus*, respectively, and error bars represent standard errors. Significant differences per event were calculated with a pairwise test, and p-values were adjusted with Bonferroni correction. (Significance levels: *=0.05, **=0.01, ***=0.001, ****=0.0001)

The variability observed during the first two days (Fig. 5) may have resulted from the different lifestyles of the studied eriophyoid species [18]. *P. adalius*, as a vagrant, is adapted to the flat leaf surface of a rearing arena. In contrast, a refuge-seeking lifestyle of *P. fructiphilus*, which often involves seeking refuge in areas such as flower buds and petiole bases, may lead to less frequent feeding on the arena, as these mites spend more time searching for shelter [19,20]. Both mite species demonstrated the ability to carry RRV for more than a week. The higher variability in *P. adalius* might indicate different mechanisms of RRV retention. Comparisons can be drawn with other plant-infecting members of the *Bunyavirales* and especially orthotospoviruses (family*Tospoviridae*). It has been shown that transmission dynamics differ significantly between vector species of tomato spotted wilt orthotospovirus (TSWV), the better-studied member of the group [21]. In the case of TSWV, vector competence is influenced by virus replication in larvae and migration to salivary glands. It is unclear whether emaraviruses, similar to orthotospoviruses, require acquisition during the larval, nymphal or adult stages [22] for successful transmission and whether the ability to acquire the virus changes as mites develop [23,24].

Emaraviruses and orthotospoviruses are characterized by similar genome structures and virion architectures, leading researchers to suggest that emaraviruses might be transmitted in a persistent, propagative manner, as observed for orthotospoviruses [25]. While some studies suggest a persistent, propagative mode [13,26], others propose a semipersistent mode [27]. Our study provides evidence for the replication of RRV in *P. fructiphilus*. However, these attributes and transmission characteristics may not be consistent across different emaravirus/vector/host pathosystems.

Our current understanding of the virus-mite interactome is nascent. A knowledge gap exists concerning the intricate transmission mechanisms and molecular determinants of virus dissemination in mites [6,28]. Addressing these gaps is crucial for devising innovative, selective, and durable control measures similar to other groups of viral pathogens [29–32]. Outbreaks of known and emerging arthropod-borne diseases, such as rose rosette, are increasing in frequency and scale due to factors associated with climate change, human demographics, and globalization of trade [33,34]. Our methodology, which involves quantifying virus concentrations in individual mites, offers new insight into eriophyoid-borne diseases. The presented approach is versatile enough for further analysis and applicable to other pathosystems. This study is a step toward enhancing our understanding of virus dynamics in mites and can be used to develop practical tools to combat the threats they pose to agriculture and biodiversity.

## Supplementary Information

**Additional file 1** Primers and probes used in the experiments, sequences of targeted regions, details on quantification of RRV titer using direct RT-qPCR and TaqMan assay, and qPCR standard curves generated for multiple independent runs.

## Funding information

TZD was supported by the National Science Centre in Poland (Polonez Bis-1 grant number 2021/43/P/NZ9/03267), and IET was supported by the United States National Institute of Food and Agriculture project ARK02850 and the Arkansas Agricultural Experimental Station.

## Author contributions

T.D. and I.T. conceived, designed and conducted experiments. A.R., T.D. and I.T. analyzed the data. All participated in writing the paper and internal review. All authors have read and approved the final manuscript.

## Conflicts of interest

The authors declare that there are no conflicts of interest.

## Notes

### Competing Interest Statement

The authors have declared no competing interest.

