## Supplemental file 1 for "Quantification of rose rosette emaravirus (RRV) titers in eriophyoid mites: insights into viral dynamics and vector competency"

**Supplementary Table 1** Primers and probes used for TaqMan-based RT-qPCR detection of rose rosette emaravirus, mite reference gene and generation of standard curves

| Gene | Primer or probe name | Sequences (5'-3') | Dye | Amplicon size | Reference |
|---|---|---|---|---|---|
| *TaqMan-based RT-qPCR detection* | | | | | |
| RNA3 RRV | RRV2F | TGCTATAAGTCTCATTGGAAGAGAAA | - | 104 | Dobhal et al., 2016 |
| | RRV2R | CCTATAGCTTCATCATTCCTCTTTG | - | | |
| | RRV probe-2 | TGCTAGAGA/ZEN/CATTGGTACAACAAGCAA/3IABkFG/ | FAM | | |
| 18S rDNA | 18S-TqM-F2 | TGGGTTGCGATTCCTTTAAT | - | 107 | this study |
| | 18S-TqM-R2 | AATCATACTTCCCCCGGAAC | - | | |
| | 18S-TqM-P2 | CTCCGATCA/ZEN/TTATGATCCACCCAGC/3IABkFQ/ | FAM | | |
| *Generation of standard curves* | | | | | |
| RNA3 RRV | RNA3flanF1 | CGTATTCACAAGCTAGAGACTACTCC | - | 201 | this study |
| | RNA3flanR2 | ATTGTGCACCTCTATCAGCAGCT | - | | |
| 18S rDNA | Flan18S-TqM-F2 | GATCAGATACCGCCCTAGTTC | - | 189 | this study |
| | Flan18S-TqM-R2 | CCCTTCCGTCAATTCCTTTAAG | - | | |

**An amplicon encompassing the virus (RNA3) target region:**

CGTATTCACAAGCTAGAGACTACTCCTTTCGATGATGCTATAAGTCTCATTGGAAGAGAA
AACATATCTGAAGCATATGTTGAACTTGCTAGAGACATTGGTACAACAAGCAAATCAAAG
AGGAATGATGAAGCTATAGGCAAGTTCAAAGAACTGATCAAGAACTTTGCTCCTGCTTTA
GCTGCTGATAGAGGTGCACAAT

**An amplicon encompassing the mite rDNA (18S) target region:**

GATCAGATACCGCCCTAGTTCTAACCATAAACGTTGCCAACTAGCAATTGGGTTGCGAT
TCCTTTAATCGGAGTGTAAAAACTCCGATCATTATGATCCACCCAGCGGCTCTCGTAGG
GAAACCAAAGTGTTTGGGTTCCGGGGGAAGTATGATTGCAAAGTTGAAACTTAAAGGAA
TTGACGGAAGGG

**Quantification of RRV titer using direct RT-qPCR and TaqMan assay**

*Crushing of mites and reverse transcription (RT) reactions:*

Individual eriophyoids were crushed with a metal pin (size 000) in 5 µL of TE buffer and combined with an RT mix containing 4 µL 5 × RT buffer and 100 U Maxima Reverse Transcriptase (Thermo Fisher Scientific, catalog no. EP0741), 0.4 mM dNTPs (Invitrogen, catalog no. 18427088), 300 ng random primers (Invitrogen, catalog no. 48190-011), 10 U

RiboLock RNase Inhibitor (Thermo Fisher Scientific, catalog no. EO0381) and water to 20 µL total volume. The sample was incubated at 50 °C for 1 h followed by 10 min at 75 °C.

*qPCR reactions and cycling conditions:*

Reactions were prepared with 5 µL of cDNA, 10 µL of TaqMan™ Universal PCR Master Mix (Thermo Fisher Scientific, catalog no. 4304437), 500 nM of each RRV-specific primer, and 250 nM of the corresponding probe or 900 nM of each 18S rDNA-specific primer, and 250 nM of the corresponding probe in a final volume of 20 µL. Amplifications were carried out in a CFX96 Touch real-time PCR detection system (Bio-Rad, Hercules, CA, USA).

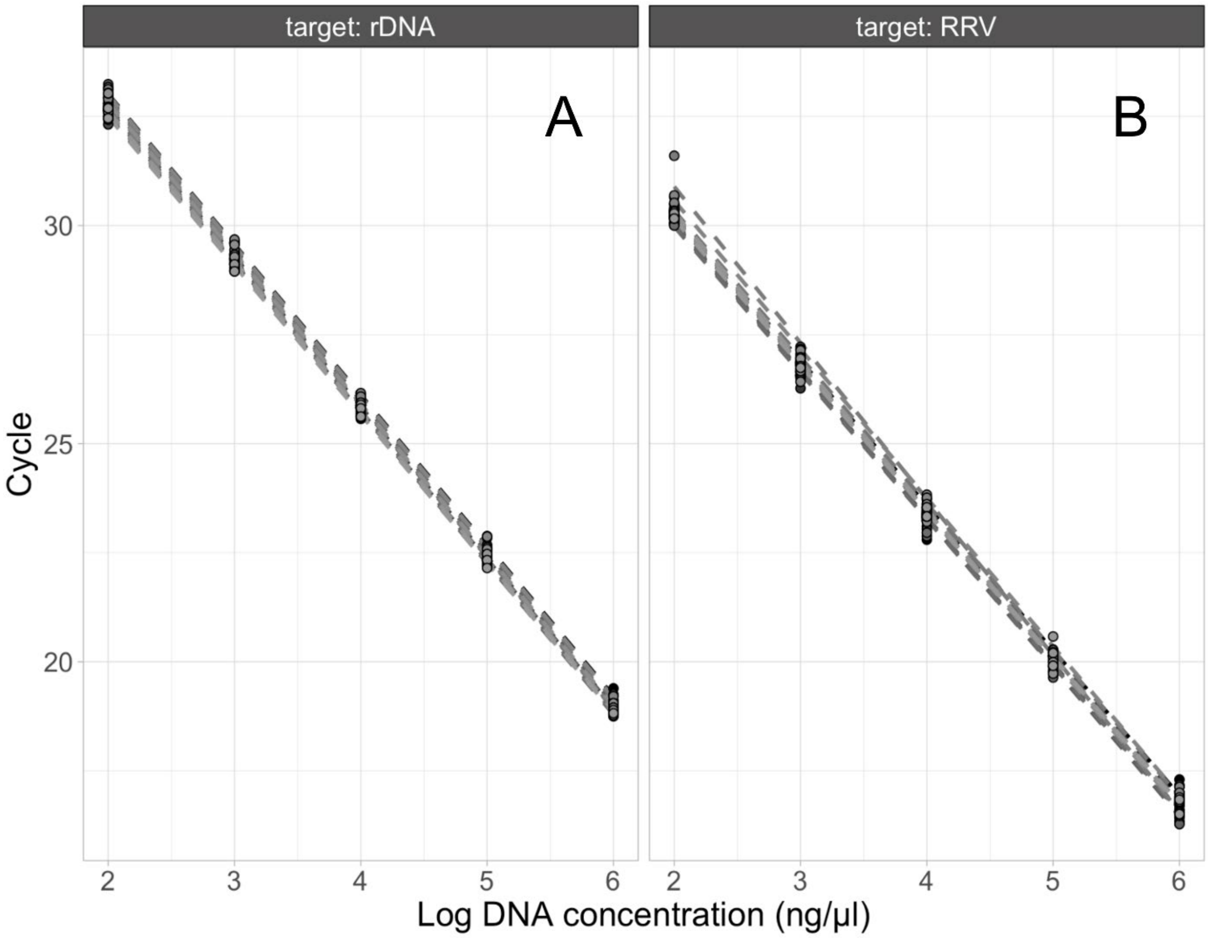


**Supplementary Fig 1** TaqMan™ qPCR standard curves generated using ten-fold dilutions of cDNA/DNA for multiple independent runs. **A**, host (mite rDNA): *Phyllocoptes adalius* and *P. fructiphilus*. **B**, viral load (cDNA). Standards were evaluated in independent assays to assess variation across plates
